# Modeling Inflammation-Driven Colon Hypertrophy and Motility Changes in Gulf War Illness

**DOI:** 10.64898/2026.01.23.701397

**Authors:** Ahilan Anantha Krishnan, Shreya A. Raghavan, Maria A. Holland

## Abstract

Gastrointestinal (GI) symptoms are a prominent feature of Gulf War Illness (GWI). Animal models attribute them to pyridostigmine bromide (PB) exposure, which induces smooth muscle hypertrophy, neuroinflammation, and motility impairment. However, animal studies only provide static snapshots of disease progression and can only partially resolve how inflammatory, neuronal, and biomechanical processes interact dynamically over time. To address this gap, we developed a computational model that couples cytokine kinetics, macrophage activation, and an excitatory-inhibitory neuronal imbalance to predict smooth muscle hypertrophy and colonic motility changes in GWI. The model was calibrated using data from mice exposed to PB under acute (7-day exposure and measurement) and chronic (7-day exposure and 30-day measurement) conditions, reproducing measured cytokine IL-6 elevations, macrophage accumulation, circular muscle thickening, and shifts in excitatory and inhibitory gene expression. Simulations captured reduced excitatory stress, and sustained loss of inhibitory relaxation, consistent with organ-bath recordings. Sensitivity analyses identified macrophage persistence as a dominant regulator of chronic inhibitory dysfunction, whereas excitatory pathways exhibited relative robustness and recovery. Thus, our model provides a systems-level view of how acute PB-induced inflammation evolves into chronic dysmotility and establishes a first step towards a virtual platform for testing hypotheses and interventions translatable to neuroimmune GI disorders.

**Highlights:**

- Neuroinflammation model predicts colon hypertrophy and motility in GWI
- Calibrated to acute (day 7) and chronic (day 30) PB-exposed mouse data
- Reproduced IL-6 rise, CD40^+^ persistence, colon thickening, and ChAT/Nos1 shifts
- Predicted excitatory stress rebound but sustained inhibitory relaxation loss

## 1 Introduction

Gulf War Illness (GWI) is a chronic condition affecting roughly one-third of the nearly 700,000 veterans deployed during the 1990-1991 Gulf War [1, 2]. Among its diverse manifestations, gastrointestinal (GI) symptoms are especially common and burdensome. Veterans report diarrhea, constipation, abdominal pain, and bloating at rates several-fold higher than those of non-deployed controls [1, 3, 4]. These symptoms often occur without overt structural lesions, suggesting that functional disturbances in the enteric nervous system (ENS) and its immune-inflammation regulation, rather than tissue injury, are the primary drivers of dysfunction. [2, 5]. A key contributor is pyridostigmine bromide (PB), administered prophylactically during deployment to prevent nerve agent toxicity. Subsequent studies have linked PB exposure to long-lasting neuroimmune changes and gastrointestinal dysfunction [1, 2, 6, 7].

Our previous work has evaluated how PB exposure alters colon function in a mouse model [4, 8]. In the acute phase, exposure to PB caused colonic smooth muscle hypertrophy, increased pro-inflammatory cytokine levels, and the recruitment of CD40^+^ macrophages to the myenteric plexus. These changes coincided with the loss of excitatory (cholinergic, ChAT) and inhibitory (nitrergic, Nos1) motor neurons, resulting in altered contractile and relaxation responses. [8]. Hernandez et al. [9] also reported reactive enteric glia, neuronal loss, and disrupted motility within days of PB exposure [9, 10]. Following up on the long-term effects after one-time PB-exposure, we found that PB induces persistent neuroinflammation and a sustained excitatory-inhibitory neuronal imbalance, resulting in chronic colonic dysmotility [4]. Hernandez et al. further showed that PB exposure leaves the enteric neuroimmune system in a primed state that appears quiescent under baseline conditions but becomes unmasked upon a secondary challenge with palmitoylethanolamide (an endogenous anti-inflammatory mediator), resulting in exaggerated cytokine responses and severe motility disturbances [11]. Together, these findings indicate that PB triggers an acute neuroimmune disruption that evolves into chronic inflammation and long-term motility impairment.

The above set of studies aligns with broader principles of inflammation-driven gut dysfunction. The ENS coordinates motility through balanced excitatory and inhibitory circuits that are highly sensitive to immune mediators [5]. Cytokines such as IL-4 and IL-13 can directly increase smooth muscle tone and alter neurotransmission, producing hypercontractility and loss of coordinated motility [12, 13]. Muscularis macrophages, which support neuro-immune homeostasis under baseline conditions, can become neurotoxic when skewed toward a pro-inflammatory phenotype, causing enteric neuron loss and impaired motility [14–16]. Similar patterns appear in conditions like irritable bowel syndrome (IBS), where transient insults leave low-grade inflammation and altered neural control that drive persistent functional symptoms [17, 18]. Although the ENS has some regenerative capacity via resident progenitors [19–21], an inflamed microenvironment can inhibit neurogenesis and repair, potentially explaining why PB-exposed tissue fails to recover normal function over time [10, 20, 22, 23]. These experimental observations suggest a convergent mechanism in GWI: immune activation, neuronal injury, and impaired regeneration interact to produce chronic dysmotility.

A key limitation of experimental approaches is that they capture only a few discrete time points—typically the end of exposure or selected post-exposure intervals—leaving the continuous evolution of inflammation, hypertrophy, and neuronal dysfunction unexplained [24]. Most studies also focus on individual pathological features in relative isolation, e.g., prioritizing inflammation, neuronal loss, or muscle hypertrophy, without capturing how these processes interact dynamically to generate chronic dysmotility. Even when multiple features are examined within the same study, as in our acute and chronic PB-exposure mouse models [4, 8], the results still provide only static snapshots of complex interactions. Computational models can overcome these limitations by capturing the *continuous* trajectory of disease and explicitly coupling interacting elements, thereby bridging cellular changes with tissue-level structural and motility changes. This approach offers a systems-level framework for understanding how PB exposure leads to persistent dysfunction in colon motility in GWI.

In this paper, we develop a computational model that integrates inflammatory signaling, neuronal changes, and smooth-muscle response to reconstruct the time evolution of PB-induced colonic dysfunction. Calibrated to discrete time points, the model predicts the time evolution of cytokines, macrophages, smooth muscle hypertrophy, and excitatory–inhibitory neuronal imbalance, as well as the corresponding contractile and relaxation stress deficits. We further use the model to generate independent exposure-duration predictions and perform sensitivity analyzes that identify the dominant regulators of chronic inhibitory dysfunction.

## 2 Methods

Here, we focus on the computational framework and summarize the experimental findings that directly inform the modeling assumptions. All experimental protocols and measurement techniques are detailed in our prior publications [4, 8]; a visual overview is provided in Figure 1. The computational approach reproduces the experimental workflow, as shown in Figure 1C. To link inflammation, hypertrophy, and active stress generation, the simulation involves three sequential steps that mimic in vivo conditions and experimental protocols.

**Fig. 1.**
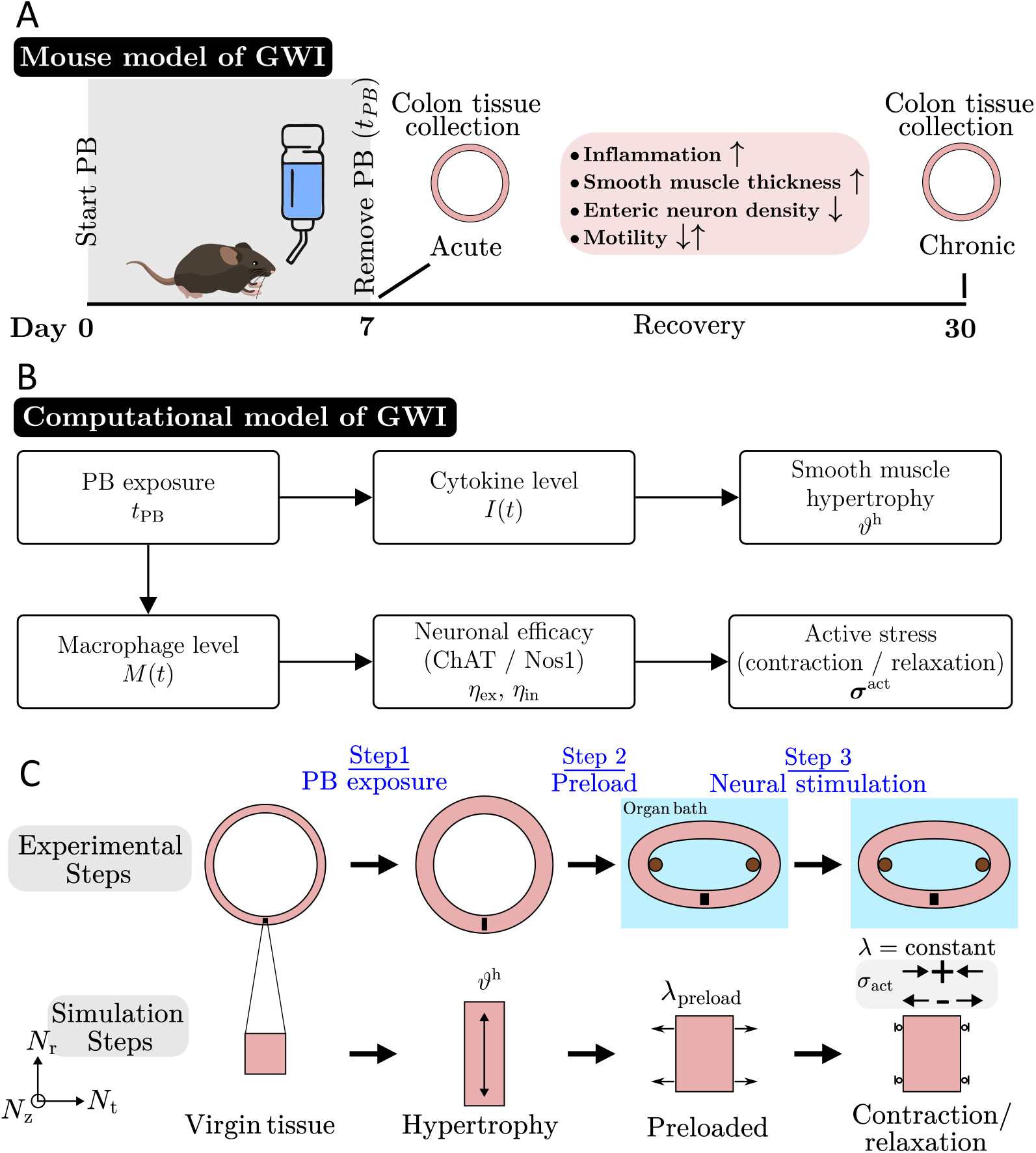
Overview of experimental and computational framework. (A) Mouse model: animals were exposed to PB for 7 days (*t*_PB_ = 7) and colonic tissue was analyzed at acute (day 7) and chronic (day 30) phases. Results showed that PB exposure induces inflammation, smooth muscle hypertrophy, enteric neuronal loss, and impaired motility, with persistent neuroinflammation and altered excitatory–inhibitory signaling in the chronic stage [4, 8] (B) Computational model: the model integrates the above findings by linking PB exposure to macrophage recruitment, cytokine production, smooth muscle hypertrophy, neuronal gene-expression changes, and disruption of contractile and relaxation responses. Together, the experimental and modeling approaches provide a systems-level view of how acute injury evolves into chronic colonic dysmotility. (C) Experimental and computational workflow. Top: schematic of the organ-bath protocol showing Step 1, PB-induced hypertrophy; Step 2, elastic preload (20%) to reproduce baseline tension; and Step 3, neural stimulation under isometric conditions. Bottom: corresponding modeling steps.

### Step 1: Inflammation-driven hypertrophy

During PB exposure, elevated cytokine levels trigger smooth-muscle hypertrophy. In the model, this process is represented through the multiplicative decomposition of the deformation gradient [25], ***F*** = ***F*** ^e^***F*** ^h^, where ***F*** is the total deformation gradient mapping the reference configuration to the current configuration, ***F*** ^h^ captures the time-dependent hypertrophy, and ***F*** ^e^ is the elastic part.

### Step 2: Elastic prestretch (organ-bath preload)

The hypertrophied tissue geometry from Step 1 serves as the new reference configuration (using ^*^IMPORT in Abaqus Standard). A purely elastic circumferential stretch *λ*_preload_ is applied to reproduce the baseline state of organ-bath preparations.

### Step 3: Isometric neural activation

The prestretched configuration from Step 2 serves as the new reference configuration for Step 3. With the total stretch fixed, excitatory and inhibitory activations are applied to simulate cholinergic contraction and nitrergic relaxation under acetylcholine and electrical field stimulation (EFS) protocols. The resulting active-stress changes under isometric conditions enable direct comparison with organ-bath recordings.

### 2.1 Kinematics

We represent inflammation-driven hypertrophy (Step 1) using the multiplicative decomposition of the deformation gradient ***F*** = ***F*** ^e^***F*** ^h^. During Steps 2–3, hypertrophy is frozen (***F*** ^h^ = ***I***), so the total deformation equals the elastic component (***F*** = ***F*** ^e^). The corresponding elastic measures are

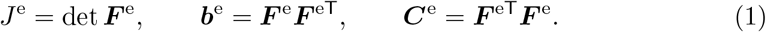

Let {***N***_r_, ***N***_t_, ***N***_z_} denote an orthonormal triad in the reference configuration (radial, tangential/circumferential, and longitudinal/axial) representing the local muscle directions. The corresponding push-forward (current) directions are defined as

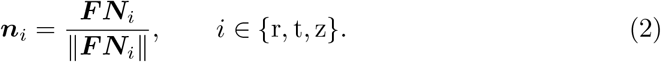

Consistent with preferential radial thickening [4], hypertrophy is prescribed as

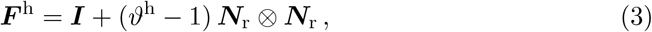

where *ϑ*^h^ is a hypertrophy multiplier that evolves as a function of the inflammatory state *I*(*t*).

### 2.2 Inflammation kinetics and hypertrophy

Inflammation *I* is quantified as the fold change in the pro-inflammatory cytokine IL-6, which rises acutely in PB-exposed colon samples [8]. For the unexposed control condition (*t*_PB_ = 0), cytokine levels remain at a physiological baseline.

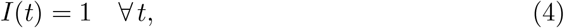

representing the absence of an inflammatory response. For PB-exposed conditions (*t*_PB_ *>* 0), the temporal evolution is modeled as

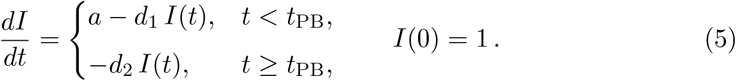

where *a* is the rise rate, *d*_1_ (set as 0 day^−1^) and *d*_2_ are the decay rates during and after exposure, respectively, and *t*_PB_ denotes the duration of PB administration, starting at *t* = 0 days. In both the acute and chronic experimental conditions, *t*_PB_ = 7 days correspond to the fixed 7-day oral PB exposure period used in our previous studies; subsequent time points (e.g., day 30) represent recovery following the end of exposure [4, 8]. This ODE-based description follows common practices in computational immunology, where cytokine levels are represented by production and decay terms [26]. The maximum attainable hypertrophy is then coupled with the time-integrated inflammation in Step 1:

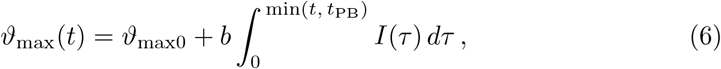

where *ϑ*_max 0_ and *b* denote the baseline maximum hypertrophy and the hypertrophy scaling coefficient, respectively, reflecting the acute circular smooth muscle thickening in our mouse models [4, 8]. The growth multiplier follows a saturating logistic-type law,

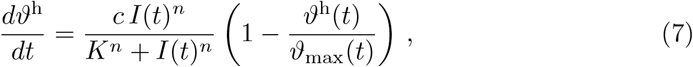

with *c* as a rate constant, *K* as a sensitivity parameter, and *n* as a Hill coefficient (see Table 1 for symbols and nominal values).

**Table 1.** Model parameters, descriptions, nominal values, and experimental data used in parameter fitting.

| Parameter | Description | Value [Units] | Experimental data used in calibration |
| --- | --- | --- | --- |
| $a$ | Inflammation activation rate | 0.9000 [day<sup>-1</sup>] | Fig. 2A |
| $d_1$ | Inflammation decay during exposure | 0.0000 [day<sup>-1</sup>] | Fig. 2A |
| $d_2$ | Inflammation decay after exposure | 0.0666 [day<sup>-1</sup>] | Fig. 2A |
| $\vartheta_{\max 0}$ | Baseline maximum hypertrophy | 6.5920 [-] | Fig. 2B |
| $b$ | Hypertrophy scaling with $\int I(\tau) d\tau$ | 0.0150 [day<sup>-1</sup>] | Fig. 2B |
| $c$ | Hypertrophy rate constant | 0.6147 [-] | Fig. 2B |
| $K$ | Hypertrophy sensitivity parameter | 0.9246 [-] | Fig. 2B |
| $n$ | Hypertrophy Hill coefficient | 4.9875 [-] | Fig. 2B |
| $m$ | CD40<sup>+</sup> activation rate | 0.0644 [day<sup>-1</sup>] | Fig. 2D |
| $y_1$ | CD40<sup>+</sup> decay during exposure | 0.0000 [-] | Fig. 2D |
| $y_2$ | CD40<sup>+</sup> decay after exposure | 0.0368 [day<sup>-1</sup>] | Fig. 2D |
| $\gamma_{\text{ch}}$ | ChAT desensitization gain | -1.0351 [-] | Fig. 4C |
| $\gamma_{\text{nos}}$ | Nos1 desensitization gain | 4.2030 [-] | Fig. 4D |
| $r_{\text{ch}}$ | Acute ChAT loss rate | 0.0942 [day<sup>-1</sup>] | Fig. 3C |
| $k_{\text{ch}}$ | ChAT recovery rate | 0.2010 [day<sup>-1</sup>] | Fig. 3C |
| $\text{ch}_{\text{ss}}$ | Steady ChAT level | 1.2334 [-] | Fig. 3C |
| $r_{\text{nos}}$ | Acute Nos1 loss rate | 0.0692 [day<sup>-1</sup>] | Fig. 3D |
| $k_{\text{nos}}$ | Nos1 adaptation rate | 0.0142 [day<sup>-1</sup>] | Fig. 3D |
| $\text{nos}_{\text{ss}}$ | Steady Nos1 level | 1.0000 [-] | Fig. 3D |
| $\mu$ | Shear modulus | 5 [kPa] | [30] |
| $\kappa_f$ | Fiber stiffness parameter | 36.0 [kPa] | Fig. 4A |
| $\alpha$ | Excitatory active stress scale | 1.7912 [kPa] | Fig. 4C |
| $\kappa_{\text{ex}}$ | Excitatory scaling rate | 0.0613 [day<sup>-1</sup>] | Fig. 4C |
| $\beta$ | Inhibitory active stress scale | 3.5531 [kPa] | Fig. 4D |
| $\kappa_{\text{in}}$ | Inhibitory scaling rate | 0.0791 [day<sup>-1</sup>] | Fig. 4D |
| $t_c$ | Chronic-case timescale | 30 [days] | [4] |

### 2.3 Constitutive equations: passive tissue response

All stresses are evaluated from the elastic state via *W* (***C***^e^, *J*^e^); in Steps 2–3, ***F*** ^h^ = ***I***, so elastic and total measures are the same. The passive response is derived from a Helmholtz free-energy density split into isotropic and anisotropic parts,

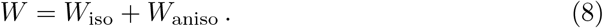

The isotropic component is a nearly incompressible logarithmic Neo-Hookean model.

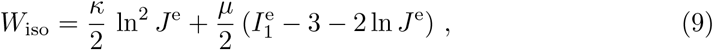

where *κ* and *µ* are the bulk and shear moduli, respectively, and the bulk modulus is set to *κ* = 100 *µ* to enforce near-incompressibility; *J* = det ***F*** = *J*^e^*J*^h^ with *J*^h^ = *ϑ*^h^, and 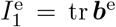. Here, *J*^e^ is the elastic volume change, and 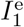 is the first invariant of the elastic left Cauchy-Green tensor ***b***^e^. The anisotropic contribution accounts for circular smooth muscle fibers via,

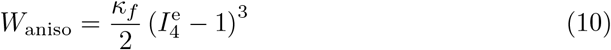

where *κ*_*f*_ is a fiber stiffness parameter, and the pseudo-invariant is defined as 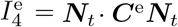. Similar anisotropic formulations have been validated in the literature for colonic tissue [27]. The passive second Piola-Kirchhoff stress is then derived using:

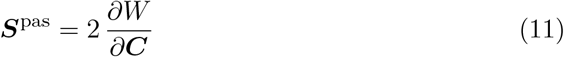

and its push-forward gives the passive Cauchy stress components as

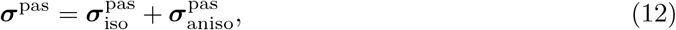

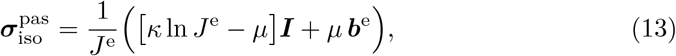

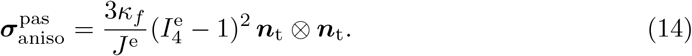

### 2.4 Active stress and neuroimmune coupling

The active response of smooth muscle fibers is regulated by neuroimmune interactions. Accordingly, the total Cauchy stress is expressed as the sum of passive and active contributions. In the acute phase of PB exposure, we observed a reduction in excitatory (ACh-evoked) contractile force, together with impaired electrical field stimulation (EFS)-induced relaxation responses [8]. During the chronic response, motility remained impaired concurrently with persistent neuroinflammation, the loss of inhibitory (Nos1^+^) neurons, and a relative increase in excitatory (ChAT^+^) neurons [4]. Based on these findings, we model the neuroimmune-mediated active stress ***σ***^act^ as a function of macrophage activation and the altered balance between excitatory and inhibitory neuronal signaling, *η*_ex_, and *η*_in_, defined as

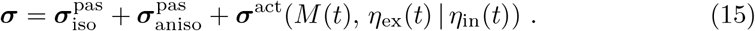

#### 2.4.1 Macrophage activation

Our prior studies demonstrated that PB exposure induces the recruitment and persistence of CD40^+^ macrophages in the myenteric plexus [4, 8]. In particular, we observed a decline in CD40^+^ signal over time in controls and an elevated, persistent trajectory following PB exposure, consistent with sustained neuroimmune signaling and smooth muscle dysfunction [28]. To capture these dynamics, we describe the relative macrophage level *M* (*t*) by

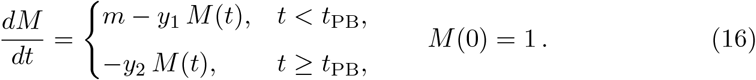

Here, *m* is the PB-driven production rate, and *y*_1_ (set as 0 day^−1^) and *y*_2_ represent the decay parameters during and after exposure, respectively. As a modeling simplification, we assume that the rates of decay in the absence of exposure (i.e., either before or after exposure) are the same, based on the observation that a gradual decline occurs in the unexposed (control) tissue as well. *M* (*t*) is normalized such that *M* (0) = 1 reflects the baseline macrophage level. Experimental CD40^+^ immunofluorescence intensities were divided by the control mean to eliminate arbitrary fluorescence units, ensuring that the model represents relative macrophage activation.

#### 2.4.2 Neuronal adaptation and desensitization

The active stress in the model consists of two components: one excitatory and one inhibitory, representing the respective contributions of cholinergic (ChAT^+^) and nitrergic (Nos1^+^) motor neurons, formulated as follows:

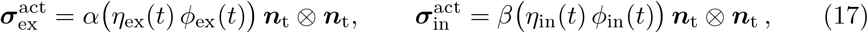

where *α* and *β* are the excitatory and inhibitory active stress scales, respectively; *η* are neuronal efficacy factors that capture both macrophage-driven desensitization and time-dependent changes in gene expression, and *ϕ* are phenomenological scaling functions. The dimensionless efficacy factors *η*_ex_(*t*) and *η*_in_(*t*) quantify the effective strength of neuronal signaling and are given by

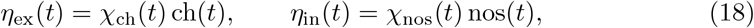

where *χ*_ch_(*t*) and *χ*_nos_(*t*) represent macrophage-mediated desensitization, and ch(*t*) and nos(*t*) denote the normalized expression levels of ChAT and Nos1, respectively.

#### Macrophage-driven desensitization

To capture the impact of persistent inflammation on neuronal responsiveness, we define exponential desensitization functions that accumulate with the integrated macrophage activation signal *M* (*t*). Here, *γ*_ch_ and *γ*_nos_ are the ChAT and Nos1 desensitization gains, respectively, and *t*_*c*_ is the chronic-case timescale:

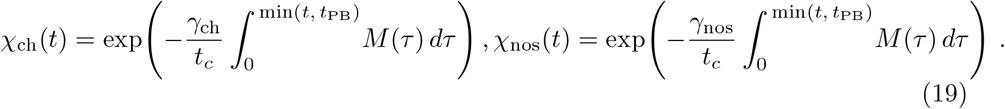

#### Neuronal loss dynamics

Exposure to PB results in early neuronal loss, thereby causing a chronic excitatory–inhibitory imbalance in the colon [4, 8]. To capture these dynamics, we describe excitatory (ChAT) and inhibitory (Nos1) gene expression as first-order processes that represent neuronal loss during PB-induced inflammation and partial recovery thereafter [4, 8]. The variables ch(*t*) and nos(*t*) denote normalized fold changes in ChAT and Nos1 expression (baseline = 1 at *t* = 0), and the PB exposure duration is *t*_PB_. Each evolves as:

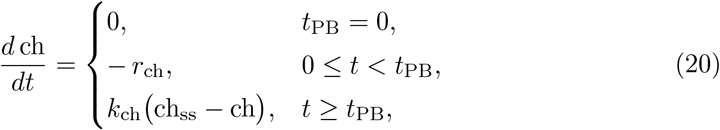

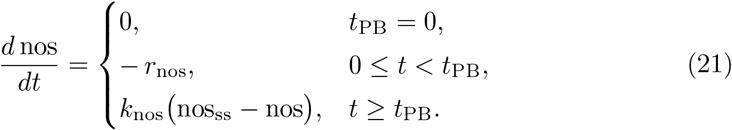

Here *r*_•_ (day^−1^) represents the acute rate of neuronal loss or transcriptional suppression during PB exposure, driven by IL-6–mediated inflammation and macrophage neurotoxicity; *k*_•_ (day^−1^) governs post-exposure adaptation once PB is withdrawn; and •_ss_ is the chronic steady-state level maintained under persistent low-grade inflammation.

#### Phenemological scaling functions

To reproduce the observed temporal evolution of neuronal function after PB exposure, we introduce phenomenological scaling functions *ϕ*_ex_(*t*) and *ϕ*_in_(*t*), which modulate the effective excitatory and inhibitory responses. These logistic type functions are not mechanistically derived; instead, they serve to capture trends observed experimentally: partial excitatory recovery and sustained inhibitory loss [4, 8]. Similar time-dependent functions have been used in computational neuroscience to represent evolving control or adaptation signals, such as in motor planning or decision-making [29]. We define them as follows:

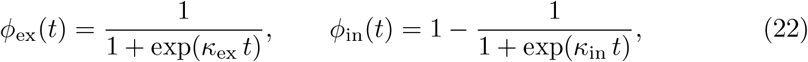

where *κ*_ex_ and *κ*_in_ are rate parameters.

### 2.5 Tangent moduli

The Lagrangian moduli are then derived by

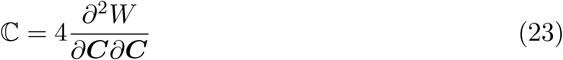

All tangent operators are derived from *W* (***C***^e^, *J*^e^). In Steps 2–3, the elastic measures are equal to their total counterparts because ***F*** ^h^ = ***I***. The push forward of the Lagrangian moduli ℂ into the spatial configuration gives the fourth-order Eulerian moduli **c**, which consist of three additive components: the isotropic material stiffness 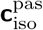, the anisotropic material stiffness 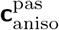, and the geometric stiffness **c**^*′*^, such that

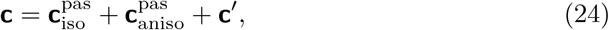

where

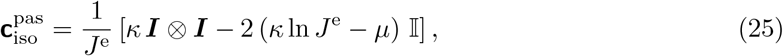

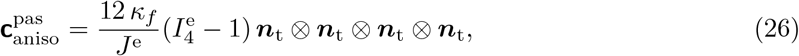

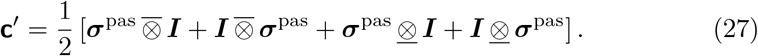

where 𝕀 is the fourth-order symmetric identity tensor, and the component representations of the non-standard fourth-order products expand as 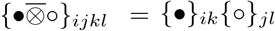 and {•<u>⊗</u>◦}_*ijkl*_ = {•}_*il*_{◦}_*jk*_.

### 2.6 Data processing for model calibration

Raw histological, biochemical, and physiological datasets from our previous studies were converted to fold-changes or dimensionless quantities to calibrate our model parameters. Circular smooth muscle thickness was measured from Hematoxylin & Eosin (H&E) images and expressed relative to the age-matched control mean to obtain the growth multiplier *ϑ*^h^. Cytokine levels were obtained from multiplex immunoassays on colon lysates (LMMP preparations) and converted to fold change *I*(*t*) with the healthy baseline set to 1. Macrophage activation was quantified as CD40^+^ immunofluorescence within the myenteric plexus and reported as the fold change *M* (*t*) after dividing each intensity by the control mean; since only day 7 and day 30 were available, we fitted the production/decay model in Eq. (16) to these time points and back-extrapolated *M* (0) to enforce the normalization *M* (0) = 1. Gene expression for ChAT and Nos1 was processed from qPCR and reported as relative expression ch(*t*) and nos(*t*) (healthy baseline = 1). Organ-bath recordings of acetylcholine-evoked contraction and EFS-evoked relaxation were converted to circumferential Cauchy stress *σ*^act^ by dividing the measured peak force *F*_max_ by the average tissue cross-sectional area, as a one-to-one geometric correspondence was not available for all specimens. The resulting values, expressed in kPa, provided consistent stress magnitudes suitable for direct comparison with model predictions. Together, these steps produced a consistent set of normalized readouts - *I*(*t*), *M* (*t*), *ϑ*^h^(*t*), ch(*t*), nos(*t*), and *σ*^act^(*t*) - used for parameter estimation and model evaluation.

### 2.7 Model calibration and evaluation

The constitutive model was implemented as a user material (`UMAT`) in Abaqus Standard, and simulations were performed using a single brick element (C3D8). All model calibrations were performed in `Python` using `NumPy` and `SciPy`. Nonlinear least-squares optimization employed `scipy.optimize.minimize` with the interior-point algorithm. Inflammation parameters (*a, d*_1_, *d*_2_) were fitted to the IL-6 fold-change trajectories *I*(*t*) for the PB-exposed group (*t*_PB_ = 7 days) at days 7 and 30. Hypertrophy parameters (*ϑ*_max 0_, *b*) were calibrated against the circular smooth muscle thickness fold-change *ϑ*^h^(*t*) using control and PB-exposed group data at day 30, while the parameters (*c, K, n*) were calibrated using data at days 7 and 30. Macrophage-kinetic parameters (*m, y*_1_, *y*_2_) were fitted using normalized CD40^+^ fluorescence intensities. Neuron gene-expression parameters (*r*_ch_, *k*_ch_, ch_ss_, *r*_nos_, *k*_nos_, nos_ss_) were fitted to the mean ChAT and Nos1 expression levels at days 7 and 30 in PB-exposed tissue. The chronic timescale *t*_*c*_ was fixed at 30 days to match the chronic observation window. The passive material parameters (*µ, κ*_*f*_) were fitted to circumferential stretch–stress data from Gong et al. [30]. Peak contractile and relaxation forces (*F*_max_) from organ bath measurements were converted to circumferential stresses by dividing each force by an estimated cross-sectional area, computed using the average smooth muscle thickness from histology at the appropriate time point and an assumed uniform sample width of 1 mm. Active-stress parameters (*α, β, κ*_ex_, *κ*_in_, *γ*_ch_, *γ*_nos_) were calibrated against organ-bath data using the combined *t*_PB_ ∈ {0, 7} conditions. After calibrating all parameters, the resulting model was used to predict responses for shorter (*t*_PB_ = 3 days) and longer (*t*_PB_ = 10 days) exposure durations. Model performance was assessed by the root-mean-square error (RMSE) between simulated and experimental values and was used as the objective function in the minimization problem. Model parameters, units, nominal values, and the experimental data used to fit the parameters are summarized in Table 1.

## 3 Results

To investigate how pyridostigmine bromide (PB) exposure drives neuroimmune disruption and colonic dysmotility, we built a computational framework around our established acute and chronic PB exposure mouse models of Gulf War Illness (GWI) (Figure 1). The framework links PB exposure to macrophage recruitment, cytokine production, smooth muscle hypertrophy, neuronal gene-expression changes, and the resulting imbalance in excitatory and inhibitory active stress generation. In this way, the model builds directly on the animal studies, complementing them by providing temporal continuity and mechanistic insights that extend beyond what can be captured at discrete experimental time points.

### 3.1 Cytokine kinetics and smooth muscle hypertrophy

Pro-inflammatory cytokine IL-6 rose sharply to 7.3-fold above baseline at day 7 under *t*_PB_ = 7 (Figure 2A). Although levels declined by day 30, they remained elevated compared to controls, indicating a sustained inflammatory response. The computational model captured this trajectory and showed that increasing *t*_PB_ amplifies and prolongs IL-6 elevation, with a smaller peak for *t*_PB_ = 3 and a more persistent response for *t*_PB_ = 10. Thus, longer PB exposures drive stronger cytokine surges and delayed resolution phases that enhance downstream tissue remodeling.

**Fig. 2.**
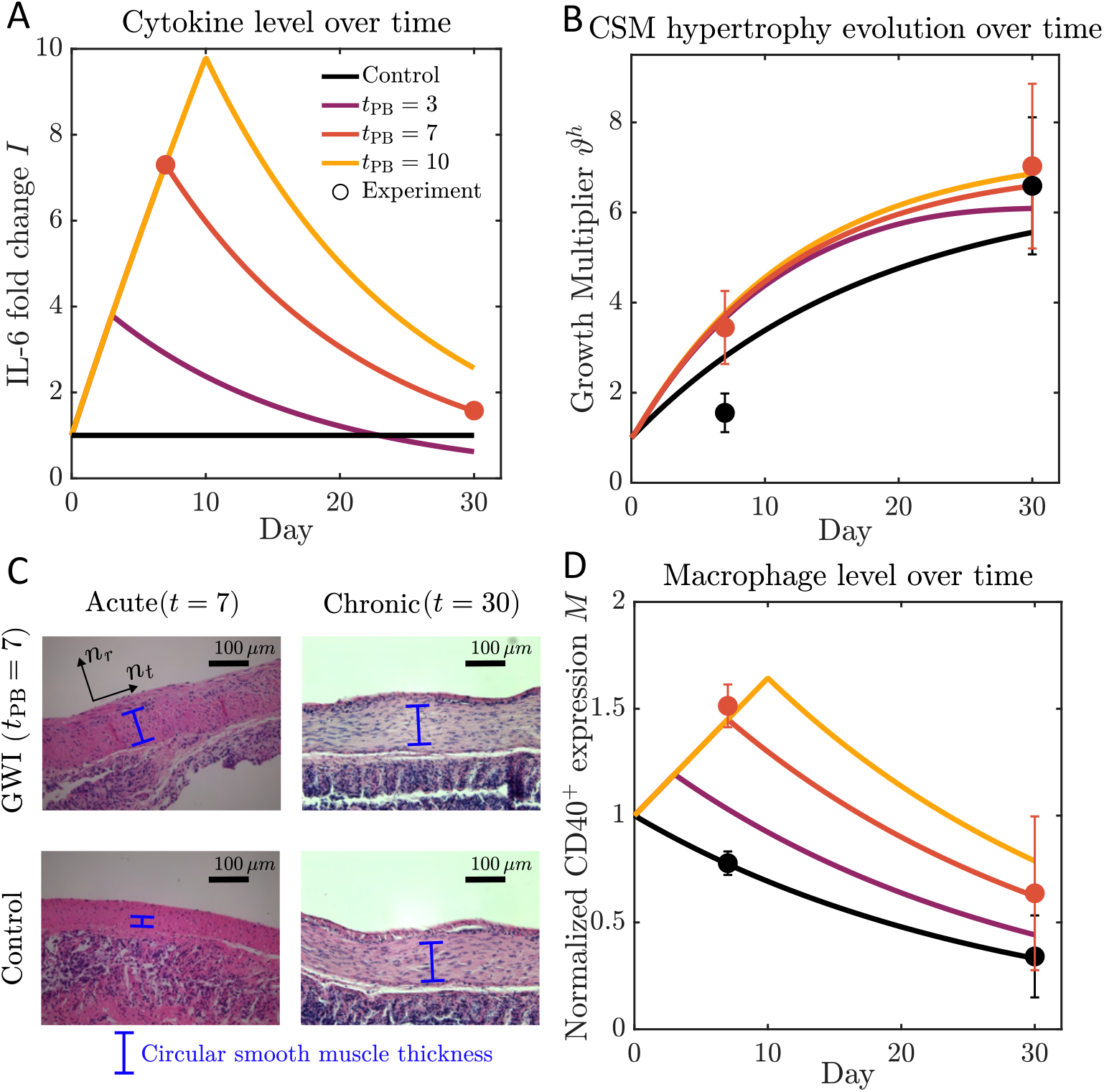
Cytokine response, macrophage activation and hypertrophy. (A) IL-6 cytokine fold-change *I*(*t*): experimental data at control and *t*_PB_ = 7 were used to calibrate the cytokine level; trajectories for *t*_PB_ = 3, 10 are model predictions. (B) Circular smooth muscle hypertrophy: Model trajectories capturing both the rapid increase during PB exposure and the sustained thickening at later stages. (C) Representative histology images of circular smooth muscle showing increased thickness in PB-exposed mouse (blue bars). (D) Normalized macrophage level *M* (*t*) (CD40^+^ fold change relative to control): experimental control and *t*_PB_ = 7 data were used for calibration; *t*_PB_ = 3, 10 represent predictions.

Muscle thickness increased 3.44-fold at day 7 and 7.02-fold by day 30 for *t*_PB_ = 7, compared with a 6.59-fold increase in age-matched controls [4, 8]. At day 7, PB exposure produced a 2.22-fold thicker circular muscle layer relative to day 7 controls (unpaired *t*-test, *p <* 0.0001), whereas the day-30 difference was not significant (1.07-fold; *p* = 0.23) (Figure 2**B**). The computational model reproduced both the rapid hypertrophy observed during PB exposure and its persistence into the chronic phase (Figure 2**B**). Notably, model trajectories showed that the magnitude of thickening was proportional to cumulative inflammatory exposure, consistent with experimental observations linking IL-6 burden to bowel wall remodeling [31, 32]. Histological analysis confirmed pronounced circular smooth muscle hypertrophy in PB-exposed tissue (Figure 2**C**).

### 3.2 Macrophage activation and neuronal gene-expression dynamics

PB exposure induced a marked increase in CD40^+^ immunofluorescence within the myenteric plexus, which we used as a surrogate measure of macrophage activation (Figure 2**D**). Normalized CD40^+^ fluorescence increased to approximately 1.5 at day 7 for the PB-exposed case relative to day 7 controls (control = 0.77; unpaired *t*-test, *p <* 0.001) [4, 8]. Although the signal declined to around 0.6 by day 30, it remained higher than the corresponding control level (0.34; unpaired *t*-test, *p <* 0.01), indicating persistent macrophage activation within the myenteric plexus. The computational model reproduced these dynamics and predicted that longer exposures would further amplify the response, with *t*_PB_ = 10 reaching nearly 1.6 during exposure. These results indicate that macrophage activation persists beyond the exposure window and provide a mechanistic basis for its role as a chronic driver of neuroimmune disruption.

In the computational framework, macrophage activation was coupled with a time-dependent desensitization function, *χ*(*t*; *t*_PB_), which modulates neuronal responsiveness during and after PB exposure. For short exposures, *χ*_ch_ (excitatory) decayed only modestly and recovered rapidly, followed by a further decline, whereas *χ*_nos_ (inhibitory) declined more steeply and remained low throughout the chronic phase (Figure 3**A,B**). With longer PB exposures, *χ*_ch_ drops sharply and then rebounds above its initial level once PB is removed. In contrast, *χ*_nos_ declines rapidly and remains low even after the removal of PB. These envelopes describe the effective temporal profiles governing neuronal adaptation and recovery.

**Fig. 3.**
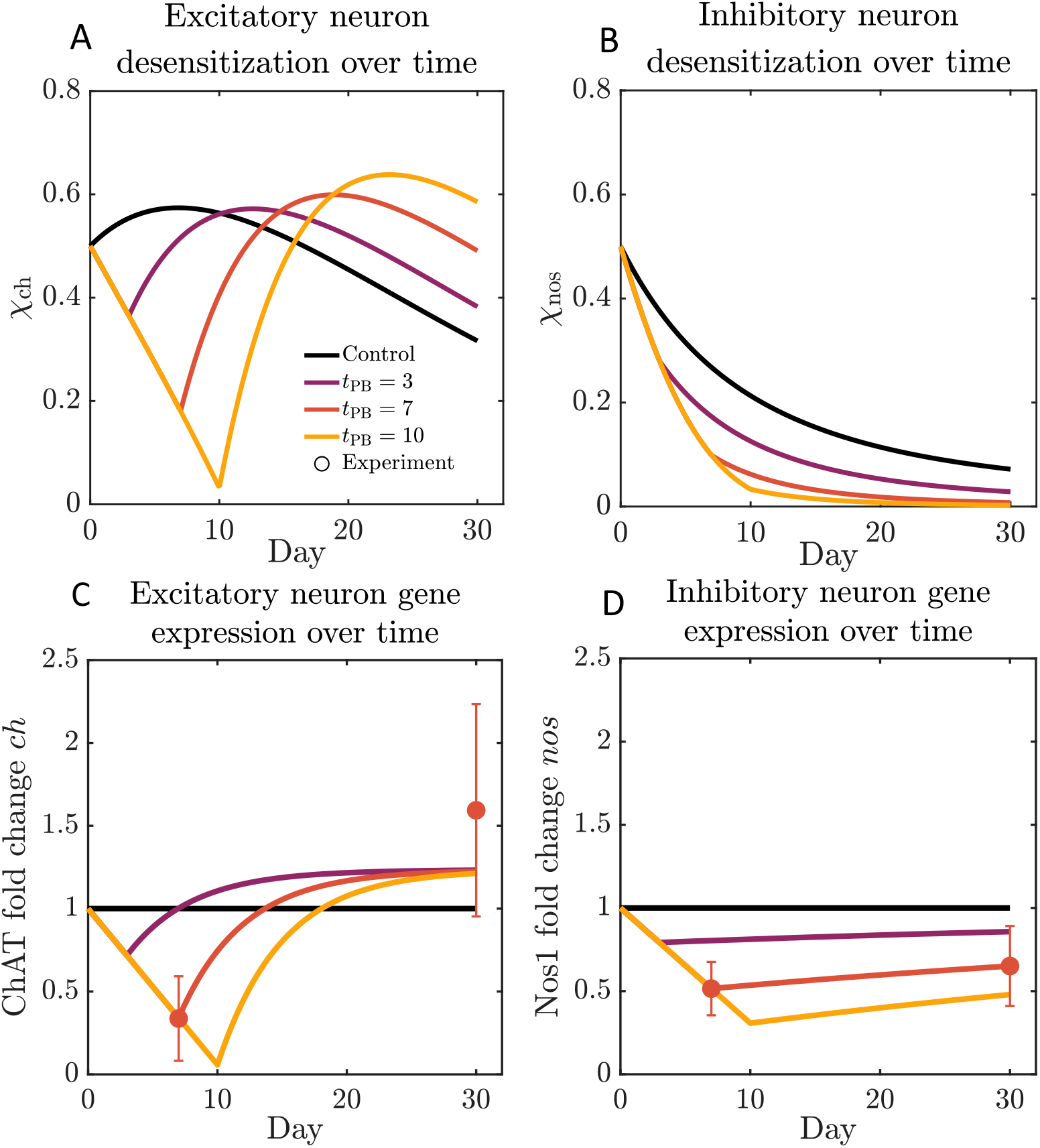
Neuronal gene-expression dynamics. (A) Excitatory neuron desensitization *χ*_ch_(*t*) under shorter exposures show partial recovery, whereas longer exposures cause sustained loss. (B) Inhibitory neuron desensitization *χ*_nos_(*t*) shows a monotonic decline with exposure duration. (C) Excitatory (ChAT) neuron gene expression dropped acutely to 0.34-fold at day 7, consistent with transient excitatory dysfunction, but reversed to above baseline by day 30 (1.59-fold). (D) Inhibitory (Nos1) neuron gene expression was more persistently reduced, with levels near 0.51-fold at day 7 and remaining suppressed (0.65-fold) by day 30. Together, these trajectories illustrate a temporal imbalance: early excitatory disruption followed by sustained inhibitory loss, a pattern captured by the computational model and aligned with experimental observations.

PB exposure also altered neuronal gene-expression profiles, which were considered a surrogate for neuron prevalence or loss. Here, the expression of ChAT and Nos1 serves as markers for excitatory and inhibitory neurons, respectively (Figure 3**C,D**). ChAT expression was acutely suppressed to 0.34-fold at day 7 but recovered above baseline by day 30 (1.59-fold), consistent with transient excitatory dysfunction followed by over-recovery. Nos1 expression declined to 0.52-fold at day 7 and remained near 0.65-fold at day 30, indicating a persistent inhibitory impairment. These changes reflect altered neuronal composition; the sustained Nos1 deficit is consistent with the partial loss of nNOS^+^ neurons, whereas ChAT recovery suggests a restoration of excitatory capacity. The computational model reproduced these trajectories, predicting similar acute suppression and partial recovery for ChAT, and sustained Nos1 reduction, with longer exposures yielding deeper and more persistent inhibitory loss.

### 3.3 Contractile and relaxation stress adaptation

Integration of neuronal and inflammatory modules with the active-stress component allowed the model to reproduce the observed contractile and relaxation responses (Figure 4). In calibrated simulations (*t*_PB_ = 7 days), the excitatory active stress decreased from a baseline of 0.90 kPa to 0.33 kPa at day 7 and then recovered to 0.88 kPa by day 30. Consistent with the experiments, PB-exposed colons showed a markedly reduced response compared with healthy controls at day 7 (0.14 kPa vs. 0.94 kPa); however, by day 30, the excitatory stress not only recovered toward the healthy baseline; but also exceeded the time-matched control (0.89 kPa in PB-exposed vs. 0.44 kPa in healthy tissue), indicating an overcompensatory excitatory rebound (Figure 4**C**). In addition, the independent predictions for *t*_PB_ = 3 and *t*_PB_ = 10 days revealed a suppression response during exposure, followed by a recovery that became more pronounced with longer exposure.

**Fig. 4.**
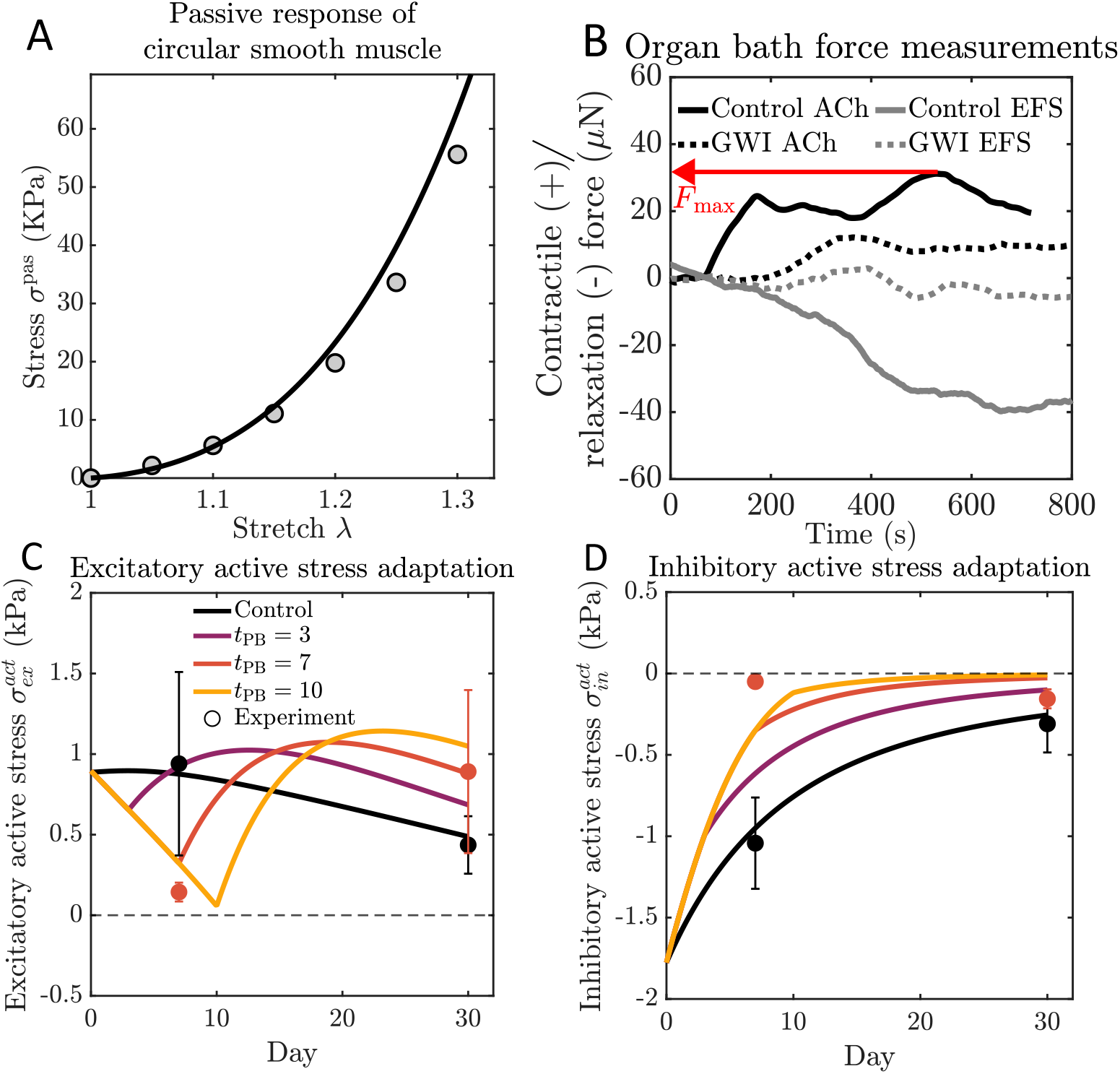
Active stress adaptation. (A) Passive stretch-stress response: the material model was tuned to capture the experimental measurements (open circles) from Gong et al. [30]. (B) Representative organ bath recordings: tissues were set to 20% baseline stretch in 37 °C DMEM+HEPES; excitatory protocol used acetylcholine (ACh, 1 µM/ mL); inhibitory responses were evoked by EFS (40 V, 10 Hz, 0.3 ms pulses for 30 s). The maximum force is each recording was noted and used to calculate the stress which was used in the subsequent model calibrations [4, 8] (C) Excitatory active stress: experimental data (*t*_PB_ = 7) showed a sharp reduction at day 7 with partial recovery by day 30. Model predictions suggest that longer exposures (*t*_PB_ = 10) would exacerbate the early drop and delay recovery. Dashed line represents zero stress reference. (D) Inhibitory active stress: baseline relaxation capacity was progressively diminished by day 30 in calibrated simulations, with stronger chronic suppression predicted at higher *t*_PB_. Together, these results link neuronal imbalance to measurable deficits in contractility and relaxation.

Experimental measurements showed a severe loss of inhibitory relaxation capacity at day 7 in PB-exposed tissue, with the inhibitory active stress reduced to −0.05 kPa compared with −1.04 kPa in healthy controls. By day 30, the inhibitory response remained markedly blunted (−0.16 kPa in PB-exposed vs. −0.31 kPa in healthy tissue).

In calibrated simulations (*t*_PB_ = 7 days), the inhibitory active stress reproduced this pronounced impairment, reducing from a baseline of −1.78 kPa to −0.05 kPa at day 7 and recovering only partially to −0.16 kPa by day 30. Unlike the excitatory pathway, which showed substantial rebound, the inhibitory pathway remained chronically suppressed, consistent with persistent deficits in nitrergic neuronal function (Figure 4**D**). The independent predictions for *t*_PB_ = 3 and *t*_PB_ = 10 days revealed progressively stronger and more persistent inhibitory suppression with increasing exposure duration.

Together, these results highlight a temporal shift from acute excitatory dysfunction to sustained inhibitory impairment, consistent with experimental observations.

### 3.4 Cytokine sensitivity analysis

Perturbing cytokine kinetics substantially altered the IL-6 trajectories (Figure 5). For *t*_PB_ = 7, the baseline parameter set produced a 7.5-fold increase in IL-6 by day 7, followed by a decline toward near-baseline by day 30. Increasing the IL-6 production rate *a* by 50% elevated the acute peak to approximately 11-fold, whereas reducing *a* lowered it to roughly 4-fold. Modulating the decay rate in the absence of exposure (*d*_2_) produced opposite effects: slower decay maintained IL-6 above fourfold at day 30, while faster decay accelerated its return to baseline. Despite these pronounced kinetic shifts, downstream hypertrophy remained relatively buffered (Figure 5**B**). Across all perturbations, hypertrophy at day 30 varied by less than 10% (ranging from 6.0 to 6.6), indicating that sustained hypertrophy is governed more by the cumulative inflammatory burden ∫*I*(*t*) *dt* than by transient IL-6 peaks.

**Fig. 5.**
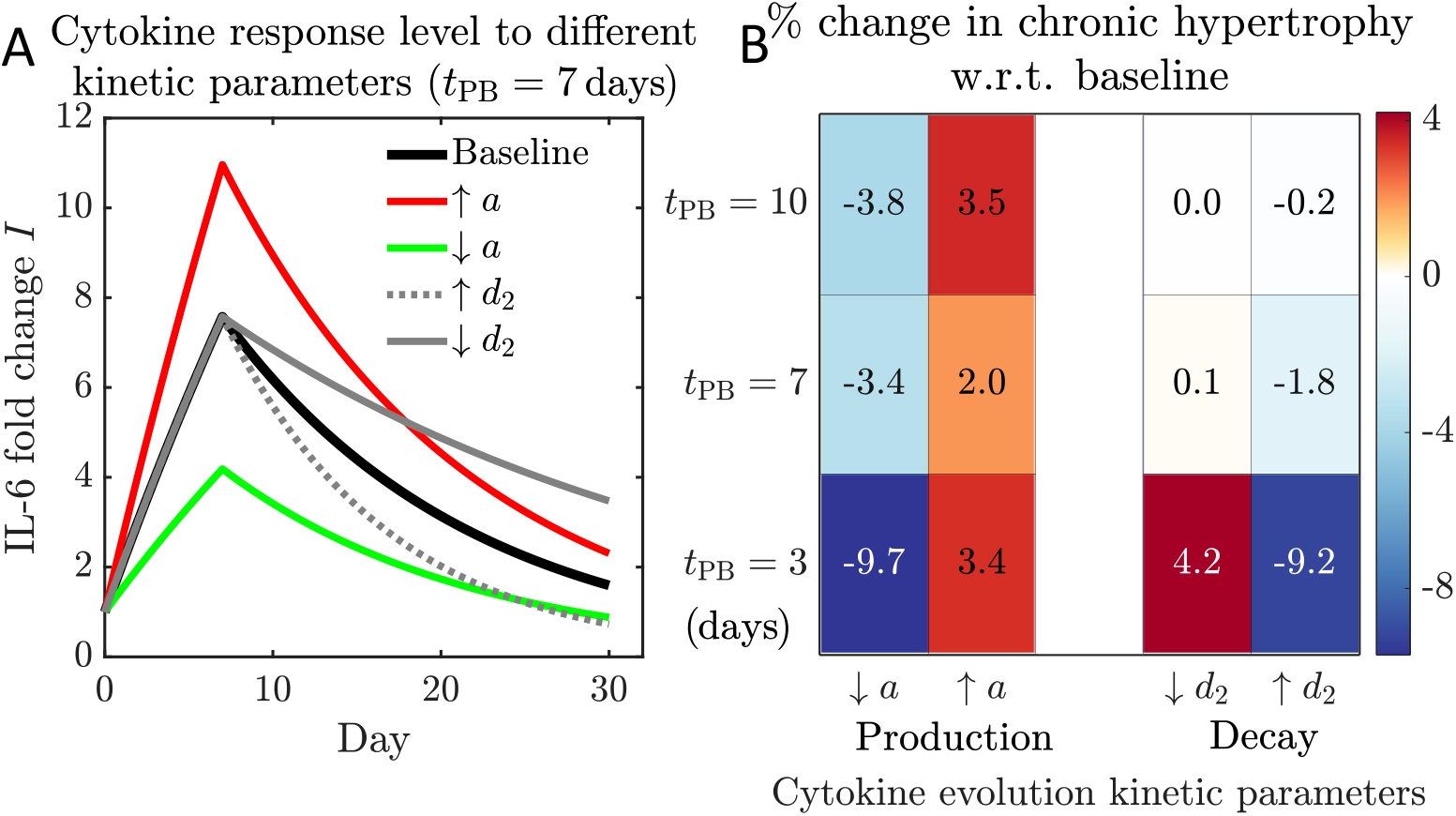
Cytokine sensitivity analysis. (A) For *t*_PB_ = 7 days, increasing the IL-6 production rate *a* amplified the acute peak, while reducing *a* lowered it; likewise, increasing the decay rate (post-exposure rate *d*_2_) accelerated recovery, whereas decreasing them prolonged IL-6 elevation. (B) Chronic hypertrophy changed only modestly across these perturbations (*<* 10%), indicating that the hypertrophy multiplier *ϑ*^h^ depends primarily on cumulative inflammatory exposure rather than transient cytokine peaks.

### 3.5 Macrophage sensitivity analysis

Macrophage kinetic parameters exert a pronounced influence on the active-stress response (Figure 6). In the acute regime, excitatory stress exhibited minimal sensitivity to changes in either the macrophage production rate *m* or the post-exposure decay constant *y*_2_ (Figure 6**B**), with variations confined to approximately ±6%. In contrast, inhibitory stress responded relatively strongly to macrophage perturbations (Figure 6**C**), spanning roughly −20% to +25% depending on whether *m* was increased or decreased. In the chronic regime (*t* = 30 days) and across all exposure durations (*t*_PB_ = 3, 7, 10 days), excitatory stress remained comparatively stable (Figure 6**D**), varying within a modest range of approximately −19% to +24% across perturbations in *m* and *y*_2_. Inhibitory stress, however, showed far greater sensitivity to macrophage kinetics (Figure 6**E**). Increasing the macrophage production rate *m* yielded the strongest suppression of inhibition (up to −57%), whereas decreasing *m* caused an increase in inhibitory stress (up to ∼ +137% compared to the baseline). Modulating macrophage clearance through *y*_2_ produced similarly large effects: faster clearance alleviated the decrease in stress (up to ∼ +67%), while slower clearance intensified it (down to ∼ −50%). Together, these results show that macrophage persistence governed by *m* and *y*_2_ is the primary driver of long-term inhibitory dysfunction, whereas excitatory output remains relatively resilient to similar inflammatory perturbations.

**Fig. 6.**
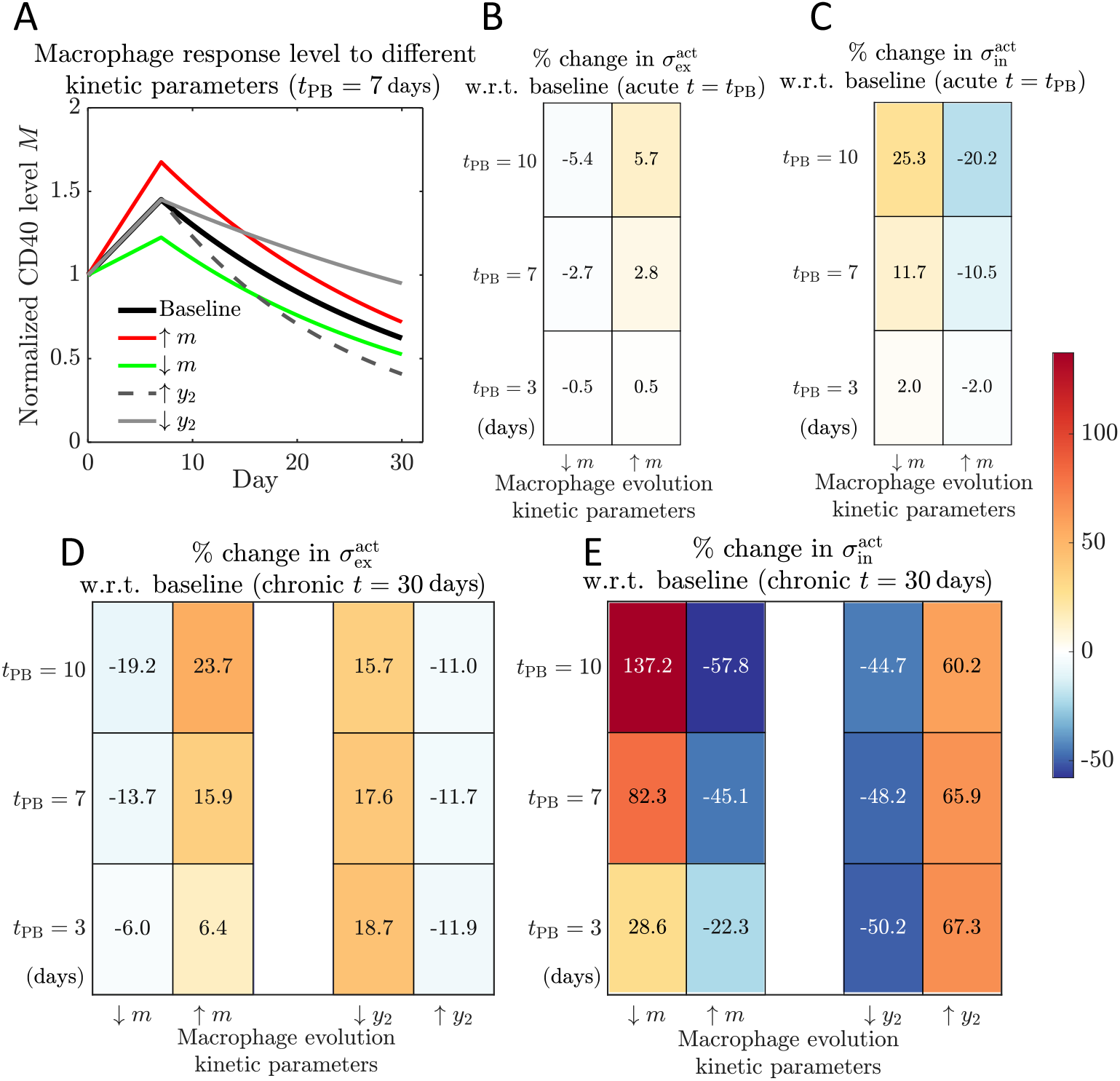
Macrophage sensitivity analysis. (A) Perturbing macrophage kinetics altered *M* (*t*) trajectories: higher production raised the peak, slower clearance prolonged elevation. (B–C) *Acute* effects: excitatory stress was stable, while inhibitory stress shifted moderately. (D–E) *Chronic* effects: excitatory stress remained robust, but inhibitory stress was highly sensitive, with reduced production or faster clearance rescuing muscle function and the opposite changes suppressing it. Percentage changes are reported relative to the baseline: 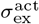 changes are computed using the signed stress, whereas 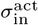 changes are computed using their absolute value. Accordingly, positive values indicate increased excitatory stress or stronger inhibitory relaxation (larger inhibitory magnitude).

## 4 Discussion

### Inflammation-induced remodeling and dysmotility

Our computational model reproduces how an acute neuroinflammatory response drives structural remodeling in the colon and subsequent motility dysfunction. PB exposure triggers a surge of pro-inflammatory cytokines (notably IL-6) and an influx of M1-polarized (CD40^+^) macrophages in the colonic neuromuscular layer, consistent with PB-treated mice [8]. These immune responses promote neuroplastic changes; for example, the loss of inhibitory nitrergic neurons is reproduced, mirroring Nos1^+^ motor neuron loss in chronic GWI models [4]. The model also captures smooth muscle hypertrophy and altered contractility arising from persistent inflammation, aligning with impaired colonic motility and muscle remodeling reported *in vivo* [4, 8]. Together, these mechanisms illustrate how inflammation-induced ENS injury produces long-term dysmotility.

### Broader context in neuroimmune related GI disorders

The model’s cascade of inflammation → ENS disruption → and motility dysfunction parallels other neuroimmune GI conditions, including IBD, IBS, and post-infectious enteropathies. In IBD, prolonged inflammation leads to neuronal loss or phenotypic changes, alongside smooth muscle hyperplasia, contributing to dysmotility [33, 34]. Even after acute inflammation resolves, colonic neuroplastic changes persist, disrupting motility [35]. Similar low-grade inflammation is implicated in IBS and functional disorders, where immune cell infiltration and cytokine elevation correlate with altered enteric neuron function and changes in bowel habits [36, 37]. A single exposure (e.g., infection or toxin) can, therefore, induce lasting gut neuroinflammation, a mechanism underlying both post-infectious IBS and GWI alike [36]. The reproduction of persistent CD40^+^ macrophage activation and cytokine elevation in our computational GWI model suggests that it could potentially be extended to represent broader neuroimmune gut dysfunction. Growing experimental evidence highlights the overlap between GWI dysmotility and other functional disorders, revealing their shared pathogenic elements, where sustained inflammation and ENS remodeling prevent the restoration of neuroimmune homeostasis [8, 35]. Taken together, our model represents the first step towards a potential computational platform for exploring how neuroimmune interactions drive long-term changes in gut motility.

### Modeling contributions and novel insights

Our computational framework complements experiments and yields new insights into gut neuroimmune dynamics. First, it provides continuous temporal dynamics linking acute exposure to chronic outcomes, whereas animal studies capture only discrete snapshots. By simulating days to weeks of cytokine-smooth muscle interactions, the model reveals how an initial inflammatory spike transitions into a sustained low-grade state with progressive tissue changes—a process that is difficult to observe empirically. Second, it enables sensitivity analyzes to identify which parameters (e.g., macrophage activation, cytokine production, neuronal death) most influence outcomes. These analyzes pinpoint critical *control knobs*, highlighting potential therapeutic targets or biomarkers. Third, *in silico* perturbations allow for testing hypotheses that are impractical to test *in vivo*—for example, inhibiting IL-6 signaling or enhancing excitatory neuron survival to predict downstream effects. Such experiments generate testable predictions and prioritize interventions, as seen in other complex disease models [24, 38]. Overall, our model integrates diverse data (immunological, neuronal, mechanical) and probes dynamic hypotheses that are difficult to assess *in vivo*, deepening our understanding of GWI pathogenesis.

### Comparison to existing GI and immunological models

This work bridges a gap by integrating neuroimmune interactions into GI modeling. Prior models of GI function have focused on either neural circuits or immune responses, rarely both. ENS models (including those with interstitial cells of Cajal as pacemakers) have simulated reflex circuits coordinating motility [38, 39], replicating peristalsis and segmentation while omitting the immune system. Conversely, immunological models of gut inflammation (ODE-based networks, agent-based models) have been used to study inflammatory bowel disease, for instance [24], but usually omit neuromuscular functions. The novelty of our model is that it explicitly links ENS remodeling and immune-driven inflammation to smooth muscle function. This neuroimmune coupling is increasingly recognized as critical for intestinal homeostasis and dysmotility [16, 40, 41]; yet, it has not been represented in prior computational studies. By including macrophage-ENS interactions, our approach offers a more comprehensive view that encompasses both motility changes and modulation by inflammation.

### Implications and use cases for experimentation and therapy

The integrated model offers practical applications as a research tool and for hypothesis generation. As an *in silico* representation of GWI colonic pathology, it could potentially serve as a virtual testbed for experiments that are impractical in vivo. For instance, researchers can simulate the timing and dose of an anti-inflammatory treatment (e.g., IL-6 inhibitor or a pro-resolving mediator) to predict whether it prevents chronic inflammation after toxin exposure. By comparing scenarios, the model could prioritize interventions for animal or clinical trials. Beyond GWI, the model’s insights could apply to general inflammatory gut diseases. Features such as macrophage-driven neuron loss, smooth muscle hyper-contractility due to low-grade inflammation, and impaired neuro-regeneration are implicated in conditions ranging from post-infectious IBS to Crohn’s strictures [34, 35]. Thus, the virtual GWI colon can probe mechanisms and interventions (e.g., vagal nerve stimulation or combinatorial cytokine blockade) that are translatable to other GI disorders marked by neuroimmune imbalance. In summary, the model complements traditional *in vivo* and *in vitro* studies, enabling rapid hypothesis testing and offering a systems-level perspective to accelerate new experiments and therapies.

### Limitations and future directions

Despite its strengths, the model has several limitations that suggest future directions. First, the immune component is simplified to a single macrophage population and a single cytokine, whereas real gut inflammation involves a complex interplay of immune system components. For instance, inflammatory T cells (Th1/Th17) and cytokines (IFN*γ*, IL-17) are central to sustaining chronic inflammation in IBD [42]. Incorporating additional cell types and pathways could capture secondary inflammatory effects, thereby improving the fidelity of the model. Second, several submodels, such as gene expression and smooth muscle desensitization, are phenomenological rather than mechanistic, constructed to match observed temporal patterns in the absence of detailed system dynamics. As the mechanistic understanding of neuroimmune interactions deepens, these formulations can be refined to reflect more biologically grounded pathways. Third, the model is zero-dimensional, neglecting spatial heterogeneity. In reality, inflammation and remodeling are patchy or localized—for instance, Crohn’s strictures develop at prior ulceration sites with distinct neuromuscular changes [34]. Introducing compartments or gradients (proximal vs. distal colon, mucosal vs. muscular layer) would enable region-specific dynamics, such as local contractility or the spread of inflammation. Finally, the model is calibrated to mouse data, and translation to humans requires caution due to species-specific differences in immune responses and ENS organization [42]. In this context, future work will integrate the computational framework with complementary *in vitro* platforms, including immune-competent 3D bioengineered colon assembloid models, to enable controlled perturbations and functional interrogation of neuroinflammation-driven colonic dysmotility [43]. Together, these extensions will enhance the model’s predictive capability and support its translational relevance to GWI and related neuroimmune GI disorders.

## Acknowledgements

A.A.K gratefully acknowledges the email communications with Dr. Sean Ward, University of Nevada, Reno, regarding smooth muscle biology; the technical discussion with Dr. Simona Carbone, Monash University, regarding neuronal electrophysiology; and numerous impromptu discussions with Dr. Rachel Branco, University of Notre Dame, on molecular neuroscience.

## Declarations

### Ethics approval and consent to participate

Not applicable

### Consent for publication

Not applicable

### Data availability

Not applicable

### Materials availability

Not applicable

### Code availability

The model developed in this paper is available at this github link: https://github.com/ahilanananthakrishnan/JMBBM GWI Model

### Author contribution

Conceptualization: M.A.H., S.A.R.; Methodology: A.A.K.; Formal Analysis: A.A.K.; Resources: M.A.H., S.A.R.; Writing - Original Draft: A.A.K.; Writing - Review & Editing: A.A.K., S.A.R., M.A.H.; Supervision: M.A.H.; Project Administration: M.A.H.; Funding Acquisition: M.A.H.

## References

[1] Steele, L.: Prevalence and patterns of gulf war illness in kansas veterans: association of symptoms with characteristics of person, place, and time of military service. American journal of epidemiology 152(10), 992–1002 (2000)

[2] White, R.F., Steele, L., O’Callaghan, J.P., Sullivan, K., Binns, J.H., Golomb, B.A., Bloom, F.E., Bunker, J.A., Crawford, F., Graves, J.C., et al.: Recent research on gulf war illness and other health problems in veterans of the 1991 gulf war: Effects of toxicant exposures during deployment. Cortex 74, 449–475 (2016)

[3] Dunphy, R.C., Bridgewater, L., Price, D.D., Robinson, M.E., Zeilman III, C.J., Verne, N.G.: Visceral and cutaneous hypersensitivity in persian gulf war veterans with chronic gastrointestinal symptoms. Pain 102(1), 79–85 (2003)

[4] Collier, C.A., Salikhova, A., Sabir, S., Raghavan, S.A.: Persistent enteric neuroinflammation chronically impairs colonic motility in a pyridostigmine bromide-induced mouse model of gulf war illness. Biology Open 14(6), 061867 (2025)

[5] Chiu, I.M., Von Hehn, C.A., Woolf, C.J.: Neurogenic inflammation and the peripheral nervous system in host defense and immunopathology. Nature neuroscience 15(8), 1063–1067 (2012)

[6] Golomb, B.A.: Acetylcholinesterase inhibitors and gulf war illnesses. Proceedings of the National Academy of Sciences 105(11), 4295–4300 (2008)

[7] Michalovicz, L.T., Kelly, K.A., Sullivan, K., O’Callaghan, J.P.: Acetylcholinesterase inhibitor exposures as an initiating factor in the development of gulf war illness, a chronic neuroimmune disorder in deployed veterans. Neuropharmacology 171, 108073 (2020)

[8] Collier, C.A., Foncerrada, S., Clevenger, A.J., Shetty, A., Raghavan, S.A.: Acute exposure to pyridostigmine bromide disrupts cholinergic myenteric neuroimmune function in mice. Advanced Biology 7(5), 2200254 (2023)

[9] Hernandez, S., Fried, D.E., Grubišić, V., McClain, J.L., Gulbransen, B.D.: Gastrointestinal neuroimmune disruption in a mouse model of gulf war illness. The FASEB Journal 33(5), 6168 (2019)

[10] Brown, I.A., McClain, J.L., Watson, R.E., Patel, B.A., Gulbransen, B.D.: Enteric glia mediate neuron death in colitis through purinergic pathways that require connexin-43 and nitric oxide. Cellular and molecular gastroenterology and hepatology 2(1), 77–91 (2016)

[11] Hernandez, S., Morales-Soto, W., Grubišić, V., Fried, D., Gulbransen, B.D.: Pyridostigmine bromide exposure creates chronic, underlying neuroimmune disruption in the gastrointestinal tract and brain that alters responses to palmitoylethanolamide in a mouse model of gulf war illness. Neuropharmacology 179, 108264 (2020)

[12] Akiho, H., Lovato, P., Deng, Y., Ceponis, P.J., Blennerhassett, P., Collins, S.M.: Interleukin-4-and-13-induced hypercontractility of human intestinal muscle cells-implication for motility changes in crohn’s disease. American Journal of Physiology-Gastrointestinal and Liver Physiology 288(4), 609–615 (2005)

[13] Chandrasekharan, B., Jeppsson, S., Pienkowski, S., Belsham, D.D., Sitaraman, S.V., Merlin, D., Kokkotou, E., Nusrat, A., Tansey, M.G., Srinivasan, S.: Tumor necrosis factor–neuropeptide y cross talk regulates inflammation, epithelial barrier functions, and colonic motility. Inflammatory bowel diseases 19(12), 2535–2546 (2013)

[14] Becker, L., Nguyen, L., Gill, J., Kulkarni, S., Pasricha, P.J., Habtezion, A.: Age-dependent shift in macrophage polarisation causes inflammation-mediated degeneration of enteric nervous system. Gut 67(5), 827–836 (2018)

[15] Muller, P.A., Koscsó, B., Rajani, G.M., Stevanovic, K., Berres, M.-L., Hashimoto, D., Mortha, A., Leboeuf, M., Li, X.-M., Mucida, D., et al.: Crosstalk between muscularis macrophages and enteric neurons regulates gastrointestinal motility. Cell 158(2), 300–313 (2014)

[16] Schneider, R., Leven, P., Mallesh, S., Breßer, M., Schneider, L., Mazzotta, E., Fadda, P., Glowka, T., Vilz, T.O., Lingohr, P., et al.: Il-1-dependent enteric gliosis guides intestinal inflammation and dysmotility and modulates macrophage function. Communications biology 5(1), 811 (2022)

[17] Bercik, P., Verdu, E.F., Foster, J.A., Macri, J., Potter, M., Huang, X., Malinowski, P., Jackson, W., Blennerhassett, P., Neufeld, K.A., et al.: Chronic gastrointestinal inflammation induces anxiety-like behavior and alters central nervous system biochemistry in mice. Gastroenterology 139(6), 2102–2112 (2010)

[18] Bercik, P., Denou, E., Collins, J., Jackson, W., Lu, J., Jury, J., Deng, Y., Blennerhassett, P., Macri, J., McCoy, K.D., et al.: The intestinal microbiota affect central levels of brain-derived neurotropic factor and behavior in mice. Gastroenterology 141(2), 599–609 (2011)

[19] Burns, A., Pachnis, V.: Development of the enteric nervous system: bringing together cells, signals and genes. Neurogastroenterology & Motility 21(2), 100–102 (2009)

[20] Burns, A.J., Thapar, N.: Neural stem cell therapies for enteric nervous system disorders. Nature reviews Gastroenterology & hepatology 11(5), 317–328 (2014)

[21] Kulkarni, S., Micci, M.-A., Leser, J., Shin, C., Tang, S.-C., Fu, Y.-Y., Liu, L., Li, Q., Saha, M., Li, C., et al.: Adult enteric nervous system in health is maintained by a dynamic balance between neuronal apoptosis and neurogenesis. Proceedings of the National Academy of Sciences 114(18), 3709–3718 (2017)

[22] Grubišić, V., McClain, J.L., Fried, D.E., Grants, I., Rajasekhar, P., Csizmadia, E., Ajijola, O.A., Watson, R.E., Poole, D.P., Robson, S.C., et al.: Enteric glia modulate macrophage phenotype and visceral sensitivity following inflammation. Cell Reports 32(10) (2020)

[23] Palanisamy, B.N., Sarkar, S., Malovic, E., Samidurai, M., Charli, A., Zenitsky, G., Jin, H., Anantharam, V., Kanthasamy, A., Kanthasamy, A.G.: Environmental neurotoxic pesticide exposure induces gut inflammation and enteric neuronal degeneration by impairing enteric glial mitochondrial function in pesticide models of parkinson’s disease: Potential relevance to gut-brain axis inflammation in parkinson’s disease pathogenesis. The international journal of biochemistry & cell biology 147, 106225 (2022)

[24] Pinton, P.: Computational models in inflammatory bowel disease. Clinical and Translational Science 15(4), 824–830 (2022)

[25] Rodriguez, E.K., Hoger, A., McCulloch, A.D.: Stress-dependent finite growth in soft elastic tissues. Journal of biomechanics 27(4), 455–467 (1994)

[26] Chow, C.C., Clermont, G., Kumar, R., Lagoa, C., Tawadrous, Z., Gallo, D., Betten, B., Bartels, J., Constantine, G., Fink, M.P., et al.: The acute inflammatory response in diverse shock states. Shock 24(1), 74–84 (2005)

[27] Fahmy, Y., Trabia, M.B., Ward, B., Gallup, L., Froehlich, M.: Development of an anisotropic hyperelastic material model for porcine colorectal tissues. Bioengineering 11(1), 64 (2024)

[28] Stakenborg, M., Abdurahiman, S., De Simone, V., Goverse, G., Stakenborg, N., Baarle, L., Wu, Q., Pirottin, D., Kim, J.-S., Chappell-Maor, L., et al.: Enteric glial cells favor accumulation of anti-inflammatory macrophages during the resolution of muscularis inflammation. Mucosal immunology 15(6), 1296–1308 (2022)

[29] Niyogi, R.K., Wong-Lin, K.: Dynamic excitatory and inhibitory gain modulation can produce flexible, robust and optimal decision-making. PLoS computational biology 9(6), 1003099 (2013)

[30] Gong, X., Xu, X., Lin, S., Cheng, Y., Tong, J., Li, Y.: Alterations in biomechanical properties and microstructure of colon wall in early-stage experimental colitis. Experimental and Therapeutic Medicine 14(2), 995–1000 (2017)

[31] Curciarello, R., Docena, G.H., MacDonald, T.T.: The role of cytokines in the fibrotic responses in crohn’s disease. Frontiers in medicine 4, 126 (2017)

[32] Li, C., Iness, A., Yoon, J., Grider, J.R., Murthy, K.S., Kellum, J.M., Kuemmerle, J.F.: Noncanonical stat3 activation regulates excess tgf-β1 and collagen i expression in muscle of stricturing crohn’s disease. The Journal of Immunology 194(7), 3422–3431 (2015)

[33] Chow, A.K., Gulbransen, B.D.: Potential roles of enteric glia in bridging neuroimmune communication in the gut. American Journal of Physiology-Gastrointestinal and Liver Physiology 312(2), 145–152 (2017)

[34] Chen, W., Lu, C., Hirota, C., Iacucci, M., Ghosh, S., Gui, X.: Smooth muscle hyperplasia/hypertrophy is the most prominent histological change in crohn’s fibrostenosing bowel strictures: a semiquantitative analysis by using a novel histological grading scheme. Journal of Crohn’s and Colitis 11(1), 92–104 (2017)

[35] Mawe, G.M., et al.: Colitis-induced neuroplasticity disrupts motility in the inflamed and post-inflamed colon. The Journal of clinical investigation 125(3), 949–955 (2015)

[36] Spear, E.T., Mawe, G.M.: Enteric neuroplasticity and dysmotility in inflammatory disease: key players and possible therapeutic targets. American Journal of Physiology-Gastrointestinal and Liver Physiology 317(6), 853–861 (2019)

[37] Lakhan, S.E., Kirchgessner, A.: Neuroinflammation in inflammatory bowel disease. Journal of neuroinflammation 7(1), 37 (2010)

[38] Chambers, J.D., Thomas, E.A., Bornstein, J.C.: Mathematical modelling of enteric neural motor patterns. Clinical and Experimental Pharmacology and Physiology 41(3), 155–164 (2014)

[39] Chambers, J.D., Bornstein, J.C., Thomas, E.A.: Insights into mechanisms of intestinal segmentation in guinea pigs: a combined computational modeling and in vitro study. American Journal of Physiology-Gastrointestinal and Liver Physiology 295(3), 534–541 (2008)

[40] De Schepper, S., Stakenborg, N., Matteoli, G., Verheijden, S., Boeckxstaens, G.E.: Muscularis macrophages: key players in intestinal homeostasis and disease. Cellular immunology 330, 142–150 (2018)

[41] Stoffels, B., Hupa, K.J., Snoek, S.A., Bree, S., Stein, K., Schwandt, T., Vilz, T.O., Lysson, M., Van’t Veer, C., Kummer, M.P., et al.: Postoperative ileus involves interleukin-1 receptor signaling in enteric glia. Gastroenterology 146(1), 176–187 (2014)

[42] Alhendi, A., Naser, S.A.: The dual role of interleukin-6 in crohn’s disease pathophysiology. Frontiers in Immunology 14, 1295230 (2023)

[43] Collier, C.A., Salikhova, A., Rengarajan, S., Tharakesh, A., Srinivasan, S., Raghavan, S.A.: Immune-competent 3d bioengineered colons for functional interrogation of neuroinflammation-induced colonic dysmotility. bioRxiv, 2025–08 (2025)

